# Biochemical evidence of furin specificity and potential for phospho-regulation at Spike protein S1/S2 cleavage site in SARS-CoV2 but not in SARS-CoV1 or MERS-CoV

**DOI:** 10.1101/2020.06.23.166900

**Authors:** Mihkel Örd, Ilona Faustova, Mart Loog

## Abstract

The Spike protein of the novel coronavirus SARS-CoV2 contains an insertion ^680^S<u>PRRA</u>R↓SV^687^ forming a cleavage motif RxxR for furin-like enzymes at the boundary of S1/S2 subunits. Cleavage at S1/S2 is important for efficient viral entry into target cells. The insertion is absent in other CoV-s of the same clade, including SARS-CoV1 that caused the 2003 outbreak. However, an analogous insertion was present in the Spike protein of the more distant Middle East Respiratory Syndrome coronavirus MERS-CoV. We show that a crucial third arginine at the left middle position, comprising a motif R**R**xR is required for furin recognition *in vitro*, while the general motif RxxR in common with MERS-CoV is not sufficient for cleavage. Further, we describe a surprising finding that the two serines at the edges of the insert **S**PRRAR↓**S**V can be efficiently phosphorylated by proline-directed and basophilic protein kinases. Both phosphorylations switch off furin’s ability to cleave the site. Although phosphoregulation of secreted proteins is still poorly understood, further studies, supported by a recent report of ten *in vivo* phosphorylated sites in the Spike protein of SARS-CoV2, could potentially uncover important novel regulatory mechanisms for SARS-CoV2.

## Introduction

The novel SARS-CoV2 coronavirus that has caused hundreds of thousands of deaths worldwide in 2020 has puzzled scientists with its high infectivity and severe consequences for infected organs (Huang et al., 2020; Sanche et al., 2020). One of the hypotheses that emerged immediately after the release of the SARS-CoV2 sequence in January (Chan et al., 2020) was that a unique insertion in its Spike protein, predicted to carry a crucial cleavage site for furin protease, could be the key for enhancing zoonotic transmission and infectivity (Andersen et al., 2020; Cheng et al., 2019; Coutard et al., 2020; Nao et al., 2017).

The coronavirus Spike glycoprotein mediates the entry of the coronavirus into the host cell (Shang et al., 2020). It is composed of two subunits (**Fig. 1A**): S1, which binds the angiotensin-converting enzyme 2 (ACE2) on the host cell surface (Hoffmann et al., 2020a; Walls et al., 2020), and S2, which mediates membrane fusion (Du et al., 2009; Lu et al., 2014). Proteolytic cleavage of the Spike protein at the S1/S2 site is essential for the infection, as the cleavage separates two functions of Spike (Heald-Sargent and Gallagher, 2012). First, the S1 fragment will bind to the ACE2 receptor, and secondly, the S2 fragment will expose hydrophobic side chains to fuse with the membrane (Shang et al., 2020).

**Figure 1.**
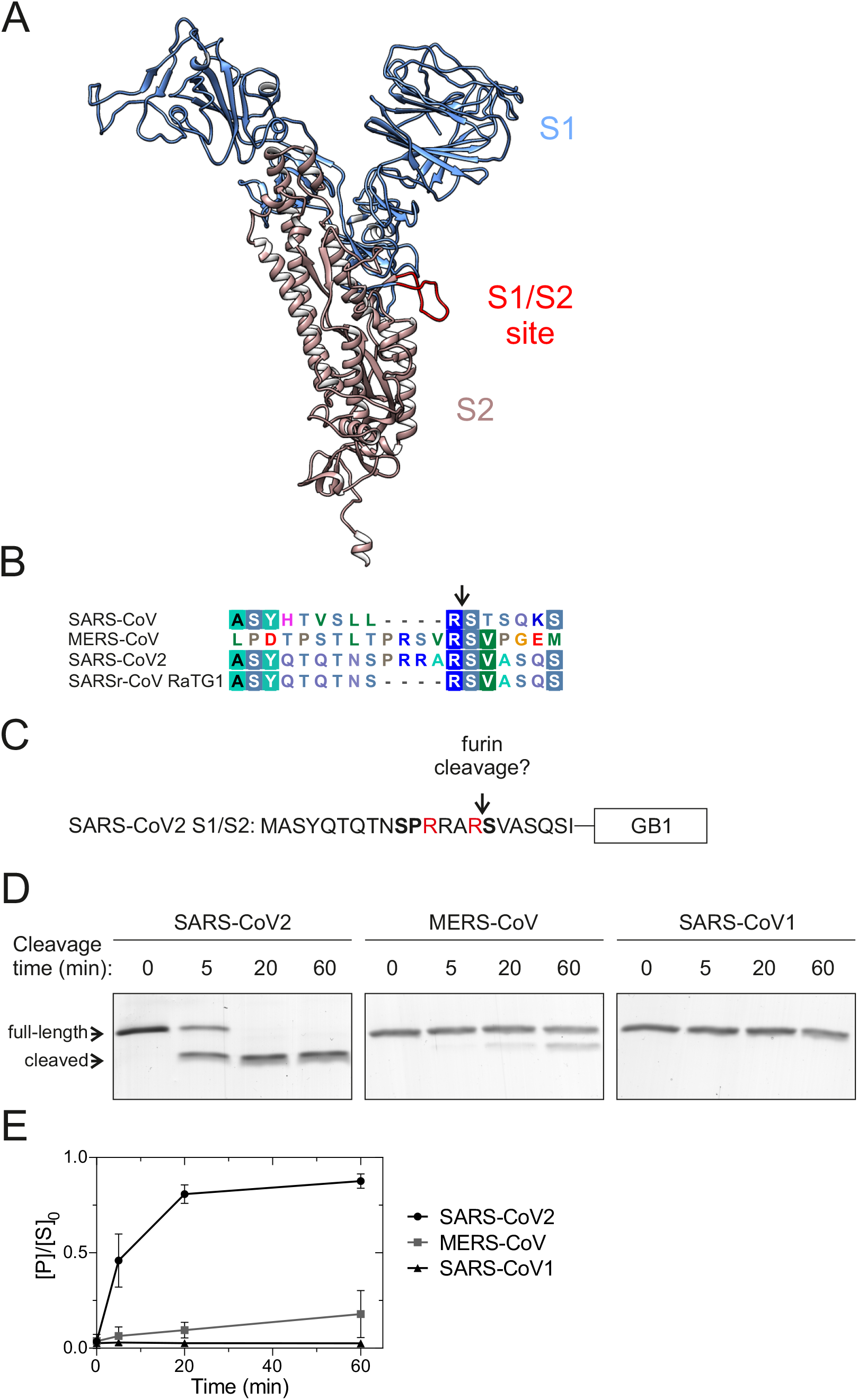
An insertion to the S1/S2 proteolytic cleavage site of SARS-CoV2 Spike protein introduces a furin site. (A) A structural model of SARS-CoV2 Spike protein (Zhang et al., 2020). Spike protein S1 (residue 1-685) is colored blue, Spike protein S2 (residue 686-1273) is colored brown and the intrinsically disordered S1/S2 proteolytic cleavage site is shown in red. The structure lacks the C-terminal residues 1148-1273. (B) Sequence alignment of the S1/S2 region in SARS-CoV, MERS-CoV, SARS-CoV2, and bat virus SARSr-CoV RaTG1, which is closely related to SARS-CoV2 — — —(Zhou et al., 2020). (C) Scheme of the constructs used to examine furin specificity. A 20-residue segment around the S1/S2 site was fused with a linker ELQGGGGG to the Streptococcal protein G B1 domain (GB1) and a C-terminal 6xHis tag. (D) Coomassie-stained SDS-PAGE gels showing the proteolytic cleavage of the GB1-fused reporter constructs. Furin activity towards the S1/S2 region of SARS-CoV2, MERS-CoV, and SARS-CoV1 was measured *in vitro* using the GB1 reporter constructs shown in ‘C’. The M_w_ of SARS-CoV2 GB1 reporter protein is 10.9 kDa, which is cleaved by furin to 9.2 kDa and 1.7 kDa fragments. (E) Quantified data from the furin cleavage assay. The plot shows the relative amount of cleaved product compared to amount of uncleaved substrate at t=0 min. Error bars show standard deviation.

The 4-residue insertion underlined in the sequence ^674^YQTQTNS<u>PRRA</u>R↓SVASQ^690^ of the SARS-CoV2 Spike protein, where the arrow indicates the S1/S2 proteolytic cleavage site, contains a previously established cleavage motif RxxR↓x for furin (Andersen et al., 2020; Walls et al., 2020) (**Fig. 1B**). After the release of the SARS-CoV2 sequence, it was also promptly realized that an analogous insertion was present in the Middle East Respiratory Syndrome coronavirus MERS-CoV that caused an outbreak in 2012 (Coutard et al., 2020). Strikingly, however, such an insertion was missing in viruses belonging to the same clade as SARS-CoV2, including SARS-CoV1, the virus causing an outbreak in 2003 (Coutard et al., 2020).

Importantly, the mechanism of proteolytic activation of Spike has been shown to play a major role in the selection of host species and the infectivity of coronaviruses (Menachery et al., 2019; Park et al., 2016). Recently, it was reported that the cleavage of the SARS-CoV2 Spike at S1/S2 is required for viral entry into human lung cells (Hoffmann et al., 2020b). However, the SARS-CoV2 spread also depends on cellular protease TMPRSS2 (Hoffmann et al., 2020b, 2020a) and the direct role of cellular furin has remained undefined. It is also not yet known if the novel sequence with the RxxR motif inserted at the S1/S2 site enables furin specificity and efficient furin-dependent cleavage. Furthermore, it has been reported that the MERS-CoV Spike, although also harboring an RxxR motif, is not activated directly by cellular furin during viral entry (Matsuyama et al., 2018). The question of Spike activation is extremely important to solve, since the initial mechanistic trigger of the current SARS-CoV2 pandemic could depend on the Spike cleavage site sequence as its host protease specificity can determine the zoonotic potential and „host-jump” of coronaviruses (Menachery et al., 2019; Park et al., 2016).

Recent Cryo-EM structures of the SARS-CoV2 Spike protein revealed that the insertion with the furin-like cleavage site emerges as a disordered loop on the side of the protein (Walls et al., 2020; Wrapp et al., 2020). Such intrinsically disordered regions (IDRs) are easily accessible for enzymes and other protein signaling modules, and often encode short linear motifs (SLiMs) that act as recognition sequences for such binding partners. Intriguingly, besides the RxxR motif, the sequence of the loop **S**<u>PRRA</u>R↓**S**V contains two serines that match the consensus of proline-directed kinases (SP), and basophilic protein kinases (RxxS), the two largest subfamilies of mammalian kinases (Hanks and Hunter, 1995; Miller and Turk, 2018). However, the Spike protein, except for its short C-terminal tail, is considered to be facing the endoplasmic reticulum (ER) or the Golgi lumen during the viral replication cycle (Lontok et al., 2004) and is not present in cytoplasm or the nucleus, where most of the kinase signaling is taking place. Nevertheless, despite still being a poorly studied field, it is well known that protein phosphorylation does not only occur on cytoplasmic and nuclear proteins, but also takes place on secreted proteins in ER and Golgi lumen, and also in the extracellular space (Dartt et al., 1996; Klement and Medzihradszky, 2017; Preisinger et al., 2004; Yalak and Vogel, 2012). Strikingly, supporting evidence for the idea of Spike phospho-regulation arises from a recent report that confirmed more than ten *in vivo* phosphorylated sites in the Spike protein of SARS-CoV2 (Davidson et al 2020).

In this study, we analyzed the furin cleavage site specificity of the SARS-CoV2 Spike using biochemical assays with substrate constructs based on the S1/S2 cleavage site sequence. We discovered that a motif RRxR, with a crucial P3 arginine (positions counted towards N terminus from the cleavage site, P4-P3-P2-P1), was required for cleavage *in vitro*, while the motif in common with MERS-CoV (RxxR) was not sufficient for furin cleavage. Further, we describe a surprising finding that the two serines flanking the insert **S**PRRAR↓**S**V can be phosphorylated by different protein kinases and both of these phosphorylations affect the ability of furin to cleave the site. Finally, we discuss the possible novel regulatory mechanisms that such interplay of three different and mutually interdependent enzymatic modifications within the insert may present for SARS-CoV2 function.

## Results

### The furin cleavage consensus site is present in the Spike protein of SARS-CoV2 but not of MERS-CoV

As the furin cleavage site is in a disordered flexible loop at the side of the Spike protein (Walls et al., 2020; Wrapp et al., 2020) (**Fig. 1A**), we set out to analyze the amino acid sequence specificity of furin cleavage using purified protein fragments corresponding to the disordered region and containing the S1/S2 cleavage site. The constructs were designed based on ^672^ASYQTQTNS<u>PRRA</u>R↓SVASQSI^692^ amino acids of Spike followed by a linker and a GB1-6xHis tag for purification (**Fig. 1C**). First, we created a set of substrate constructs based on the S1/S2 sequences of SARS-CoV2, MERS-CoV, and SARS-CoV1 (**Fig. 1B**). We followed the cleavage of the constructs by purified furin preparation in SDS-PAGE. Strikingly, we found that although the insertion in MERS-CoV has been considered as a furin cleavage site (Kleine-Weber et al., 2018; Millet and Whittaker, 2014; Park et al., 2016), it was cleaved by furin at very low efficiency (**Fig. 1D, E**). Contrarily, the novel coronavirus SARS-CoV2 site was cleaved very efficiently (**Fig. 1D, E**). Expectedly, the SARS-CoV1 motif lacking the furin site insertion showed no cleavage (**Fig. 1D, E**).

### P3 arginine in R<u>R</u>xR motif is necessary for efficient cleavage of SARS-CoV2 and MERS-CoV S1/S2 sites by furin

Next, we analyzed the furin cleavage of selected S1/S2 site mutants to better understand the specificity determinants of these sites. We used a recently described S1/S2 site mutant that lacks the four residue insertion (fur/mut (Walls et al., 2020)) (**Fig. 2A**) as a control to confirm the cleavage specificity in the *in vitro* furin assay. Deletion of the furin site in fur/mut-GB1 reporter protein abolished the proteolytic cleavage of the reporter protein (**Fig. 2B**). Next, a patient-derived mutation R682Q in Spike (ZJU-1, (Yao et al., 2020)), that changes the furin core motif RxxR↓x to QxxR↓x, also completely inhibits furin activity towards the reporter protein (**Fig. 2B**). One difference between SARS-CoV2 and MERS-CoV S1/S2 sites is the presence of arginine 3 residues upstream of the cleavage site in the former (**Fig. 2A**). Interestingly, a mutation of this P3 residue to alanine (RRAR to RAAR) greatly decreased the furin cleavage rate compared to the wild-type SARS-CoV2 sequence (**Fig. 2B, C**). Introduction of arginine to the −3 position of MERS-CoV S1/S2 (RSVR to RRVR) considerably enhanced the cleavage rate by furin, resulting in only slightly lower cleavage efficiency compared to the wildtype SARS-CoV2 S1/S2 site (**Fig. 2B, C**). These findings highlight the presence of additional specificity determinants in the proteolytic cleavage sites on top of the commonly suggested RxxR, as a furin cleavage consensus motif. Further, this result shows how a single amino acid substitution can change protease specificity, which could have impacts on the tissue tropism and host range (Millet and Whittaker, 2015).

**Figure 2.**
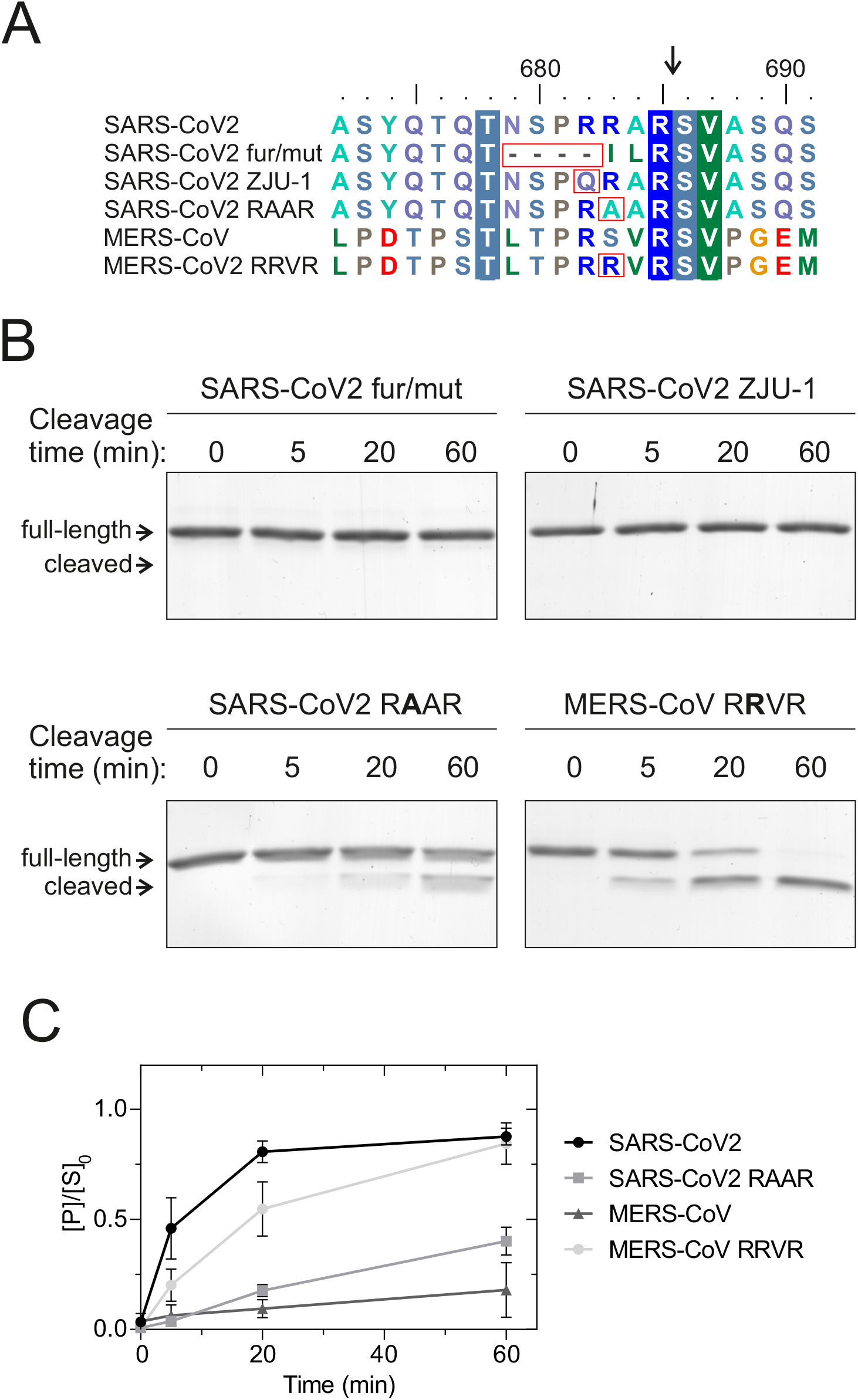
Analysis of furin cleavage efficiency of different SARS-CoV2 and MERS-CoV S1/S2 cleavage site mutants. (A) Alignment of SARS-CoV2 and MERS-CoV S1/S2 regions with the mutants used in substrate constructs for analysis of furin cleavage specificity in panel ‘B’. (B) The furin cleavage efficiency of different mutants was analyzed in an *in vitro* time-course experiment. The proteolytic cleavage of S1/S2-GB1 reporter constructs by furin is visualized by Coomassie staining of SDS-PAGE gels. (C) The plot shows the accumulation of cleaved S1/S2-GB1 reporter protein in an *in vitro* furin activity assay. The error bars show standard deviation.

### The SARS-CoV2 S1/S2 cleavage motif could be phosphorylated by proline-directed and basophilic kinases

In addition to the furin cleavage site, the four amino acid insertion to the S1/S2 site of the SARS-CoV2 Spike protein also introduces potential phosphorylation sites that flank the core furin motif (**Fig. 3A**). Spike S680 matches the consensus of proline-directed kinases (SP) and S686 forms a consensus for basophilic kinases (RxxS), two large subfamilies of mammalian kinases (Hanks and Hunter, 1995; Miller and Turk, 2018). Interestingly, the presence of potential phosphorylation sites can also be seen in the polybasic proteolytic cleavage sites of several other viral envelope proteins, including the ones of H5N1 influenza virus (**Fig. 3B**). Importantly, while the phosphorylation of Spike proteins has not been studied thoroughly, several phosphorylated residues including both SP and RxxS sites have been identified in the SARS-CoV2 Spike protein by mass spectrometry (Davidson et al., 2020).

**Figure 3.**
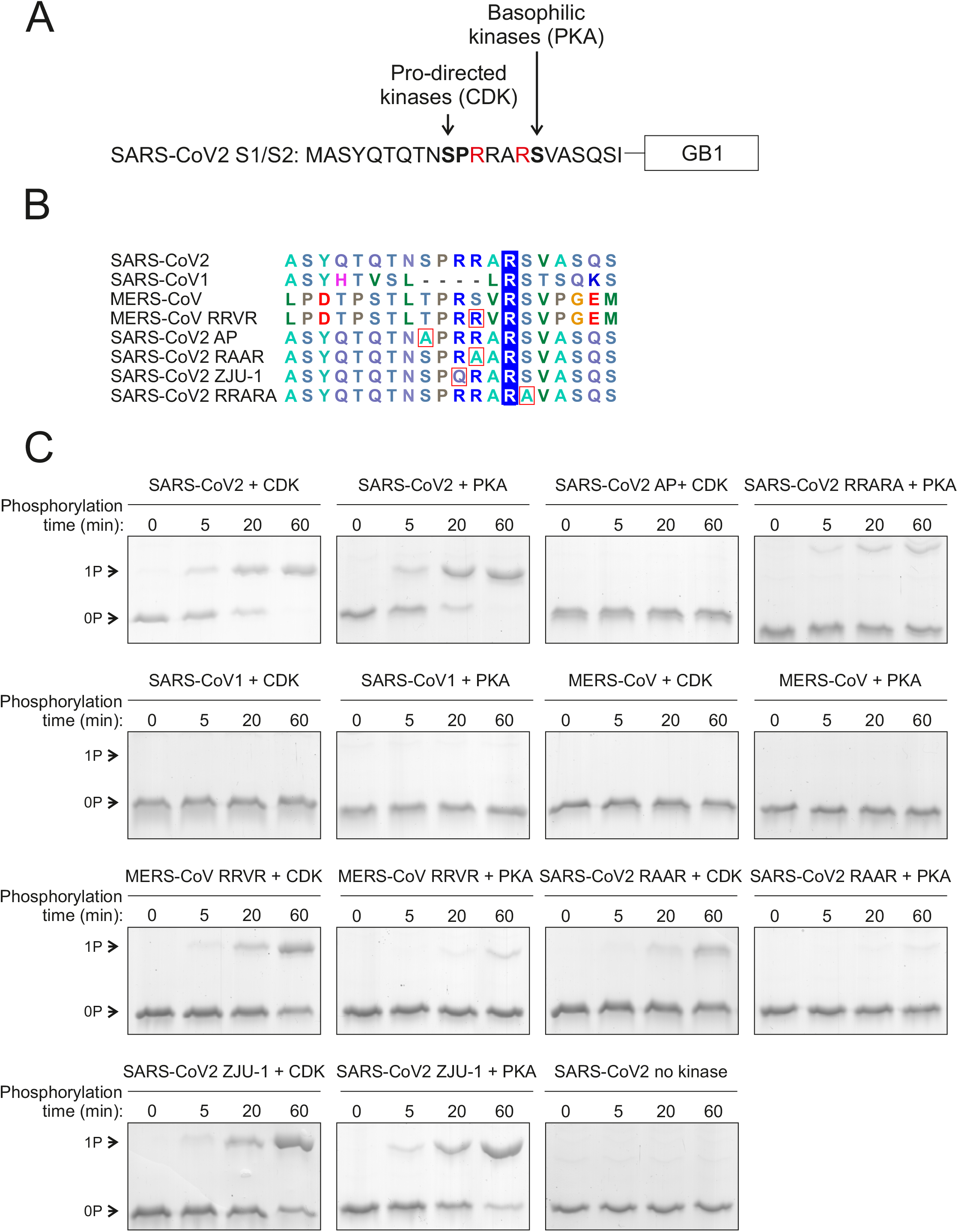
Potential phosphorylation of the S1/S2 site. (A) A scheme of the 20-residue segment of the SARS-CoV2 S1/S2 site. In addition to a furin site, the four-residue insertion (PRRA) creates potential phosphorylation sites for proline-directed kinases (S680) and basophilic kinases (S686). (B) Multiple sequence alignment of different S1/S2 sequences used in phosphorylation assays in panel ‘C’. (C) The phosphorylation of S1/S2-GB1 reporter proteins was analyzed *in vitro* using cyclin B-Cdk1 (CDK) and protein kinase A (PKA) as a representative of proline-directed and basophilic kinases, respectively. The phosphorylation reactions were stopped at 0, 5, 20, and 60 minutes. Phosphorylation shifts were analyzed using Phostag SDS-PAGE, which separates the phosphorylated form from the unphosphorylated form. Coomassie-stained gels are shown.

This prompted us to test whether the S1/S2 reporter proteins could be phosphorylated *in vitro*. For this, we used the cyclin B-Cdk1 complex (CDK) as a representative for proline-directed kinases and the protein kinase A (PKA) catalytic subunit as a representative for basophilic kinases. To examine the phosphorylation of different S1/S2-GB1 reporter proteins, we stopped the *in vitro* phosphorylation reactions at specific time points and analyzed the phosphorylation efficiency using Phos-tag SDS-PAGE to separate the phosphorylated protein from the non-phosphorylated substrate. The SARS-CoV2 S1/S2-GB1 reporter was fully phosphorylated by both CDK and PKA by the 60-minute time point (**Fig. 3C, Supplementary Fig. 1**). Mutation of the predicted phosphorylation sites, S680 for CDK and S686 for PKA, abolished or greatly reduced the phosphorylation (**Fig. 3C**). The S1/S2 segments of SARS-CoV1 and MERS-CoV were expectedly not phosphorylated by PKA (**Fig. 3C**). The SARS-CoV1 S1/S2 segment does not contain a consensus site for proline-directed kinases, and while the MERS-CoV segment contains two potential proline-directed TP sites, these sites lack a basic residue in +3 position, an important specificity determinant for CDK (Songyang et al., 1994). Introduction of the +3R (denoted as P3 arginine for furin site) that greatly increases the furin cleavage efficiency of MERS-CoV S1/S2 also enhances its phosphorylation by CDK (**Fig. 3C**). Thus, the MERS-CoV S1/S2 segment could still be phosphorylated by other proline-directed kinases. Mutations in the +2 and +3 basic residues of SARS-CoV2 S680, which were found to affect furin cleavage (**Fig. 2B**), also decrease the phosphorylation rate by CDK (**Fig. 3C**), whereas with PKA, mutation of the −3R from S686 abolishes the phosphorylation, while mutation of −4R to Q has little effect (**Fig. 3C**).

### Phosphorylation inhibits furin cleavage *in vitro*

Next, we analyzed how phosphorylation at these sites affects furin cleavage. For this, the S1/S2-GB1 reporter proteins were first phosphorylated by either CDK or PKA for 60 minutes, resulting in the phosphorylation of the majority of the substrate, followed by the addition of furin. Phosphorylation of either site adjacent to the core furin motif, S680 and S686, significantly inhibited the furin cleavage (**Fig. 4A, B**). To confirm that the effect of phosphorylation is connected to the specific site, we analyzed the cleavage of alanine mutants of the phosphorylation sites. Interestingly, alanine mutations of these sites decreased the cleavage rate, although not to the same extent as phosphorylation, and the addition of kinase to the S680A and S686A substrates did not affect their cleavage further (**Fig. 4A, C**).

**Figure 4.**
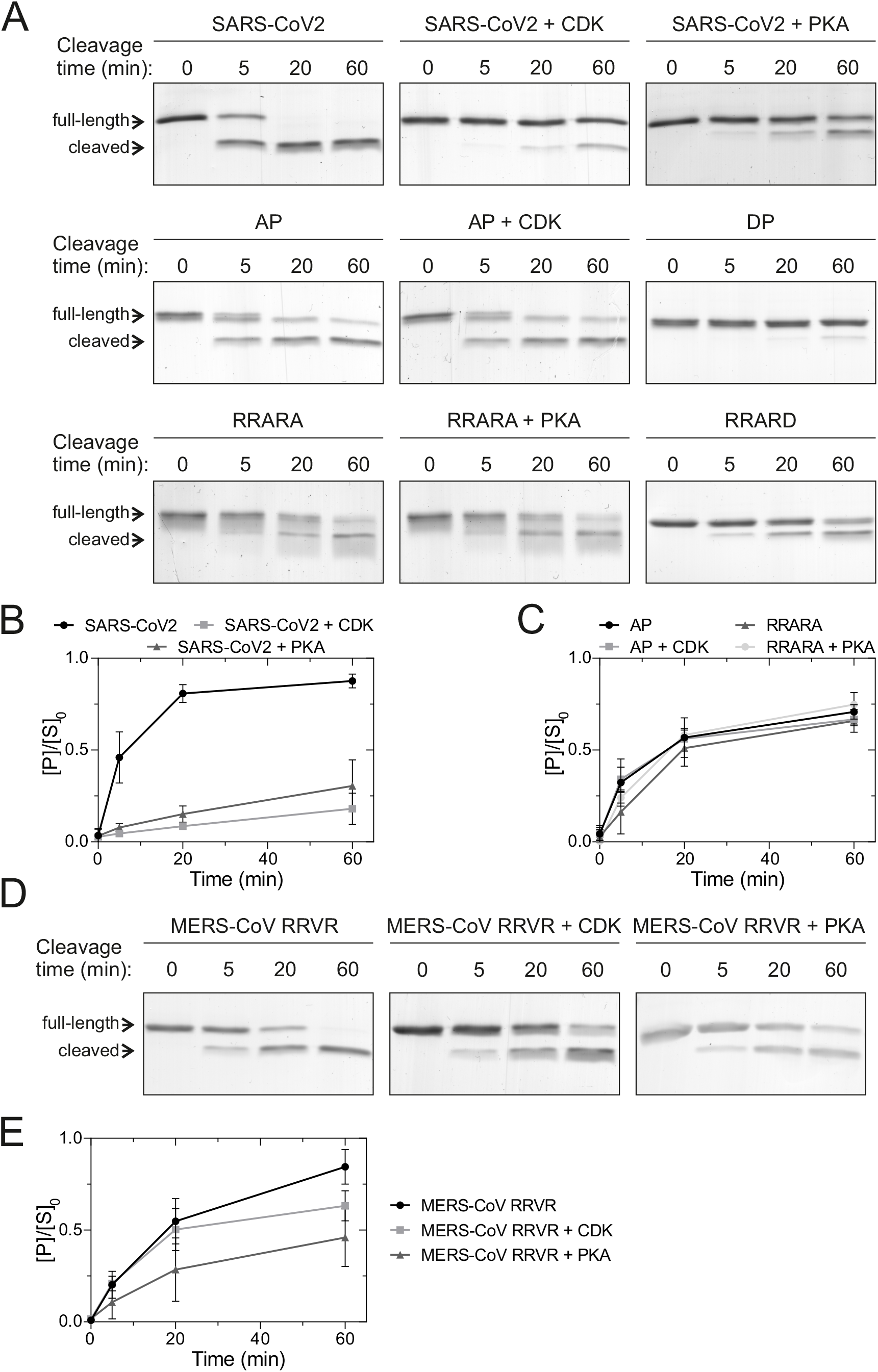
Phosphorylation at positions adjacent to the furin cleavage site inhibits the proteolytic cleavage *in vitro*. (A) The SARS-CoV2 S1/S2-GB1 reporter proteins were first phosphorylated with cyclin B-Cdk1 or PKA, followed by addition of furin. The furin activity was stopped at indicated time-points and the cleavage efficiency was analyzed by SDS-PAGE. (B) Plot showing the phosphorylation-dependent inhibition of furin activity on SARS-CoV2 S1/S2-GB1 reporter protein. Error bars show standard deviation. (C) Quantified furin cleavage profiles of the indicated SARS-CoV2 S1/S2-GB1 mutants without phosphorylation or with CDK- or PKA-mediated phosphorylation. Error bars show standard deviation. (D) Coomassie-stained SDS-PAGE gels showing the inhibitory effect of phosphorylation on furin-mediated cleavage of MERS-CoV S1/S2 with R**<u>R</u>**VR mutation. (E) Plot showing the relative abundance of cleaved MERS-CoV R**<u>R</u>**VR S1/S2-GB1 protein compared to the uncleaved form at t=0. Error bars show standard deviation.

Next, we tested the effect of mutation of the phosphorylation sites to aspartic acid, often used to mimic phosphorylation. The S680D mutation decreased furin cleavage efficiency to the same extent as phosphorylation. However, S686D had a smaller effect on furin activity, similar to S686A mutation (**Fig. 4A**). Importantly, these results show that mutations and post-translational modifications outside the core RxxR furin motif significantly affect the cleavage.

A similar phospho-inhibitory effect was seen with MERS-CoV(RRVR)-GB1 (**Fig. 4D**), although the effect was less prominent, presumably due to less efficient phosphorylation of this protein (**Fig. 3C**), resulting in incomplete phosphorylation prior to furin addition. Taken together, these data reveal that phosphorylation adjacent to the furin motif can switch off the cleavage site.

## Discussion

Our results confirm that a four amino acid insertion to the S1/S2 site of the SARS-CoV2 Spike protein introduces a furin cleavage site. However, the P3 arginine, not present in analogous insertion in MERS-CoV, is crucial for furin-dependent *in vitro* cleavage of a substrate construct carrying the sequence of the insert. While some previous reports have noted furin-mediated activation of MERS-CoV Spike (Hoffmann et al., 2020b; Kleine-Weber et al., 2018), others have argued against this due to an off-target effect of the furin inhibitor dec-RVKR-CMK (Matsuyama et al., 2018). Our finding suggests that SARS-CoV2 may have acquired a true furin cleavage specificity in contrast to the coronaviruses causing the two previous outbreaks. In addition, the discovered possibility of tight phospho-regulation at two serines creates an interesting complexity where a disordered loop carrying a short linear motif (SLiM) facilitates specific regulatory inputs for three different modifying enzymes. The *in vivo* functionality of this SLiM could emerge as one of the key functional elements in the SARS-CoV2 Spike protein, given that proteolytic processing on viral glycoproteins has been found to be a key virulence factor. Indeed, previous studies have shown that highly pathogenic influenza virus forms have been found to harbor optimal furin processing sites, whereas forms with low pathogenicity have monobasic cleavage sites (Kido et al., 2012; Sun et al., 2010). The glycoprotein cleavage specificity also determines tissue tropism, as furin is ubiquitously expressed, whereas the proteases that process monobasic cleavage sites, like TMPRSS2, are expressed mainly in the aerodigestive tract (Lukassen et al., 2020; Thomas, 2002).

The motif RxxR is often referred to as the minimal furin cleavage site, whereas RxK/RR forms an optimal motif that is cleaved efficiently (Cao et al., 2005; Krysan et al., 1999). More recent reports, however, have defined a core motif of 8 amino acids (6 residues N-terminal and 2 residues C-terminal of the cleavage site) (Tian et al., 2011). The work presented here also suggests that the furin motif is more defined and longer, as we find that RRxR is necessary for efficient cleavage of the tested S1/S2 sites by furin and that mutations flanking this core also affect the cleavage efficiency. Interestingly, the glycoprotein of highly pathogenic Ebolaviruses have been found to contain an optimal furin cleavage site, while the glycoprotein of a closely related Reston virus has a non-optimal cleavage site (Braun and Sauter, 2019; Volchkov et al., 1998). However, the requirement for P3 arginine for furin specificity discovered in this study suggests that different cellular proteases may prefer different variations of cleavage site sequences. One hypothesis would be that furin may present an activity and specificity common for different hosts and thus a window for zoonotic transfer.

Importantly, although not sufficiently studied, it is known that besides cyto- and nucleoplasm, protein phosphorylation occurs also in the extracellular space, and in lumens of ER and Golgi (Klement and Medzihradszky, 2017; Tagliabracci et al., 2013; Yalak and Vogel, 2012). For example, one of the first discovered phosphoproteins was casein, a true secreted protein (Levene and Hill, 1933). Recently, an analysis of the saliva phospho-proteome discovered close to a hundred phospho-proteins (Stone et al., 2011). Thus, despite being a secreted protein, the Spike has a potential of being regulated by protein kinases. Moreover, a recent report presented *in vivo* evidence of more than ten phosphorylated sites at the Spike protein of SARS-CoV2 (Davidson et al., 2020). Further studies are required to understand if phosphorylation of the Spike protein has a physiological role and if specific kinases are involved. One hypothesis would be that negative feedback via protein kinases could switch off the furin-dependent pre-cleavage of Spike, acting as a mitigating factor to safeguard optimal and stationary infection of cells in the tissue and preventing cell death due to overly active viral replication.

In conclusion, the described short linear motif acting as a triple regulatory module is quite unique, also in a general signaling context. Further studies are required to establish its exact role in SARS-CoV2 and also as a modular regulatory element *in vivo*.

## Methods

### Protein purification

Constructs containing a 20 amino acid region from the S1/S2 linker were fused via a linker with the sequence ELQGGGGG to the GB1 domain (immunoglobulin-binding domain of streptococcal protein G) and a C-terminal 6xHis tag. The pET28a vectors were transformed to *E. coli* BL21 cells and protein expression was induced at 37 °C by addition of 1 mM IPTG. The His-tagged proteins were purified by cobalt affinity chromatography using Chelating Sepharose (GE Healthcare). The proteins were eluted in buffer containing 25 mM Hepes-KOH (pH 7.4), 300 mM NaCl, 10% glycerol, 200 mM imidazole.

### Phosphorylation assay

The phosphorylation of the S1/S2 linker constructs was examined *in vitro* using purified cyclin B-Cdk1 (Millipore 14-450) and PKA (murine cAMP-dependent protein kinase), purified as described in (Kivi et al., 2013). The phosphorylation reactions were carried out in buffer consisting of 50 mM Hepes-KOH (pH 7.4), 150 mM NaCl, 5 mM MgCl_2_, 50 mM imidazole, 2.5% glycerol, 0.2 mg/ml bovine serum albumin, 0.15 mM EGTA, 1 mM β-mercaptoethanol, and 500 μM ATP. The kinase reactions were performed in 30 μl containing 2.5 μg of S1/S2-GB1-6xHis substrate (7.6 μM). The concentration of PKA was 375 nM and cyclin B-Cdk1 5 nM. The reactions were carried out at room temperature and were stopped at 10 s (0 min), 5 min, 20 min, and 60 min by pipetting 5.5 μl of the reaction mixture to 8 μl of 2x Laemmli SDS-PAGE sample buffer containing 1 mM MnCl_2_.

The stopped samples were incubated at 72 °C for 5 min and were loaded on Phos-tag SDS-PAGE gels containing 12.5% acrylamide, 100 μM Phos-tag, 200 μM MnCl_2_. The electrophoresis was carried out at 15 mA until the bromophenol blue dye front reached the bottom of the gel. Following electrophoresis, the gels were soaked in fixation solution (10% acetic acid, 30% ethanol aqueous solution) with gentle agitation for 15 min. The proteins were stained with colloidal Coomassie Blue G-250 (Candiano et al., 2004).

### Furin cleavage assay

The furin cleavage specificity was assayed by incubating 2.5 μg (7.6 μM) S1/S2-GB1-6xHis substrate with 2 U furin (New England Biolabs, p8077) in 30 μl reactions. The reaction buffer contained 50 mM Hepes-KOH (pH 7.4), 150 mM NaCl, 5 mM MgCl_2_, 16 mM imidazole, 2% glycerol, 0.2 mg/ml bovine serum albumin, 0.15 mM EGTA, 500 μM ATP, 0.5% Triton-X100, 2 mM CaCl_2_, 2 mM β-mercaptoethanol. In case the effect of phosphorylation on furin cleavage was studied, the S1/S2-GB1-6xHis substrate was first phosphorylated for 60 min as described above, followed by addition of furin. The reactions were carried out at room temperature and were stopped at 0, 5, 20 and 60 min by pipetting 6 ul of the reaction mixture to 2x Laemmli SDS-PAGE sample buffer.

The stopped reactions were heated at 72 °C for 5 min and loaded to 15% acrylamide SDS-PAGE. Following electrophoresis, the gels were immersed in fixation solution for 15 min and stained with colloidal Coomassie Blue G-250.

## Acknowledgments

We thank Mardo Kõivomägi, Jan Skotheim, and Dave Morgan for discussions. We are grateful to Ervin Valk, Rait Kivi and Jevgeni Mihhejev for assistance. The work was funded by the Institute of Technology basic funding grant PLTTI20909 for ML.

## Declaration of Interests

The authors declare no competing interests.

## Notes

### Competing Interest Statement

The authors have declared no competing interest.

