## Supplementarty Figure 1 for "Biochemical evidence of furin specificity and potential for phospho-regulation at Spike protein S1/S2 cleavage site in SARS-CoV2 but not in SARS-CoV1 or MERS-CoV"

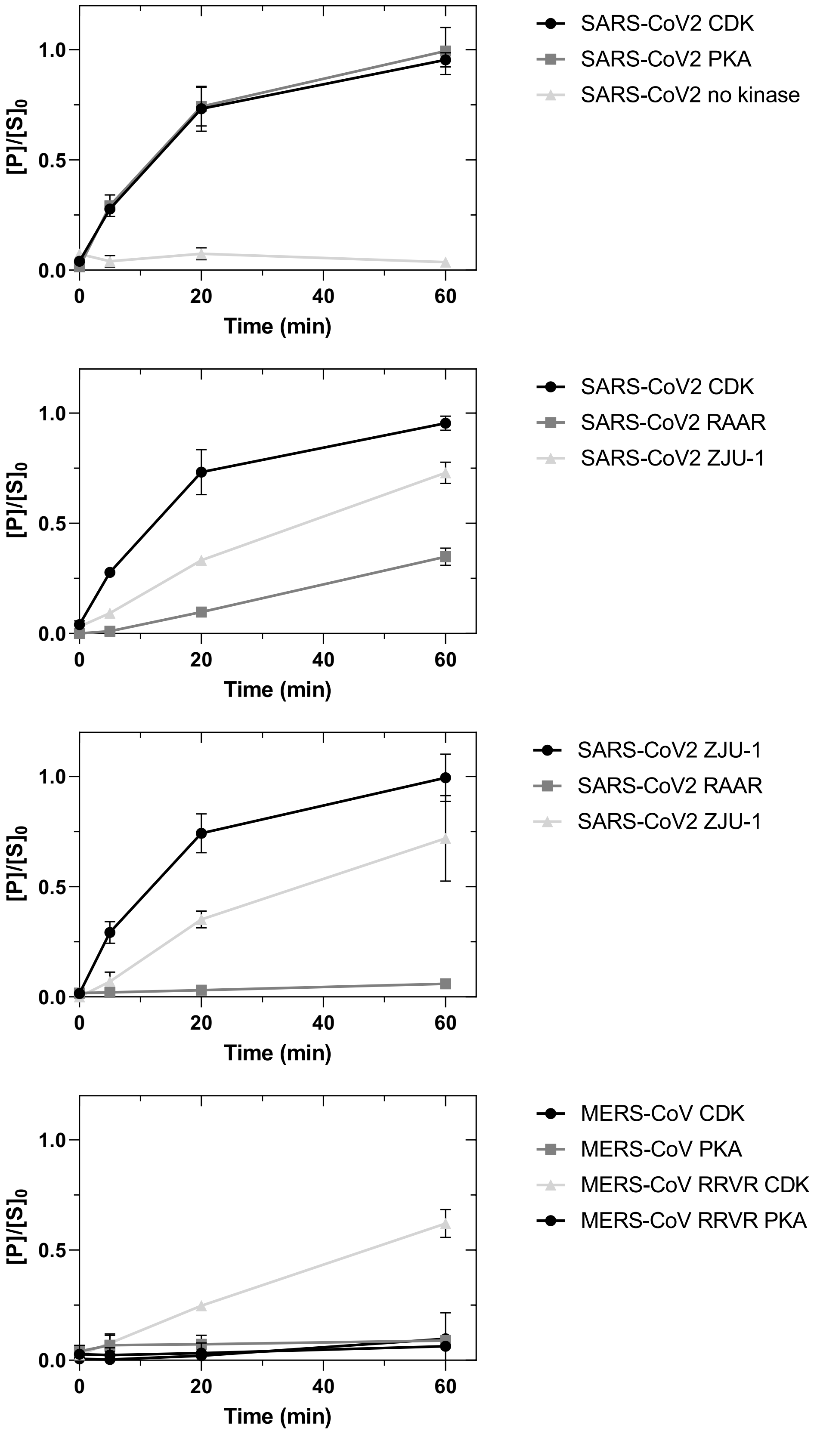


**Supplementary Figure 1. Phosphorylation of SARS-CoV2 and MERS-CoV S1/S2 sites *in vitro*.** The plots show accumulation of phosphorylated form of the indicated S1/S2-GB1 reporter protein in phosphorylation assays presented in Figure 3. The plots show relative abundance of phosphorylated form ([P]) compared to non-phosphorylated form at t=0 ($[S]_0$). Error bars show standard deviation.
